# Identification of novel mitochondrial and mitochondrial related genetic loci associated with exercise response in the Gene SMART study

**DOI:** 10.1101/2020.02.20.957340

**Authors:** NR Harvey, S Voisin, RA Lea, X Yan, MC Benton, ID Papadimitriou, M Jacques, LM Haupt, KJ Ashton, N Eynon, LR Griffiths

## Abstract

Mitochondria supply intracellular energy requirements during exercise. Specific mitochondrial haplogroups and mitochondrial genetic variants have been associated with athletic performance, and exercise responses. However, these associations were discovered using underpowered, candidate gene approaches, and consequently have not been replicated. Here, we used whole-mitochondrial genome sequencing, in conjunction with high-throughput genotyping arrays, to discover novel genetic variants associated with exercise responses in the Gene SMART (Skeletal Muscle Adaptive Response to Training) cohort (n=62 completed). We performed a Principal Component Analysis of cohort aerobic fitness measures to build composite traits and test for variants associated with exercise outcomes. None of the mitochondrial genetic variants but nine nuclear encoded variants in eight separate genes were found to be associated with exercise responses (FDR<0.05) *(rs11061368: DIABLO, rs113400963: FAM185A, rs6062129 and rs6121949: MTG2, rs7231304: AFG3L2, rs2041840: NDUFAF7, rs7085433: TIMM23, rs1063271: SPTLC2, rs2275273: ALDH18A1).* Additionally, we outline potential mechanisms by which these variants may be contributing to exercise phenotypes. Our data suggest novel nuclear-encoded SNPs and mitochondrial pathways associated with exercise response phenotypes. Future studies should focus on validating these variants across different cohorts and ethnicities.

**AUTHOR SUMMARY:** Previous exercise genetic studies contain many flaws that impede the growth in knowledge surrounding change in exercise outcomes. In particular, exercise studies looking at mtDNA variants have looked at very small portions of the mitochondrial genome. Mitochondria are the ‘power house’ of the cell and therefore understanding the mitochondrial genetics behind adaptations to training can help us fill knowledge gaps in current research. Here, we utilised a new mitochondrial genetic sequencing technique to examine all mitochondrial and mitochondrial related genetic variations. We have shown that there were no mitochondrial specific variants that influenced exercise training however there were 9 related variants that were significantly associated with exercise phenotypes. Additionally, we have shown that building composite traits increased the significance of our association testing and lead to novel findings. We will be able to understand why response to training is so varied and increase the effectiveness of exercise training on a host of metabolic disorders.

## INTRODUCTION

Responses to exercise training depends on the type of exercise stimulus, and varies considerably between individuals (1-3). This variability is tissue-specific, and may be explained by a combination of genetic variants, epigenetic signatures, other molecular and lifestyle factors (4, 5). Mitochondria are the key mediators of intracellular energy and are involved in many essential cell metabolism and homeostasis processes (6) with exercise training improving mitochondrial function and content (6-9).

The mitochondrial genome encodes 37 genes that are highly conserved but differ slightly amongst different regional isolates (haplogroups) (10). Mitochondrial haplogroups and Single Nucleotide Polymorphisms (SNPs), in conjunction with SNPs in mitochondrial-related genes (nuclear encoded mitochondrial proteins: NEMPs) have previously been associated with athletic performance in highly trained populations and response to exercise training in the general population (11). While these studies have advanced our understanding, they have primarily utilised targeted genotyping technology such as candidate gene approaches, or Sanger sequencing to investigate specific mitochondrial coding regions and NEMPs, such as *NRF2* and *PGC1α* (12-15). Many of these studies also lacked robust technical measures on aerobic fitness measures (9). As such, many of the identified variants have not been replicated, and exercise-related genetic variants remain unknown (16).

To date, studies assessing mitochondrial DNA (mtDNA) variants and NEMPs pertaining to exercise training have focused on protein-coding variants, with no studies looking at the more subtle effects of synonymous and non-coding changes (11, 17-20). Further, these studies have often based haplogroup analyses on sequencing or genotyping of the mitochondrial hypervariable region(s) (∼500-1,000bp), with no consideration for the remaining mitochondrial genome (∼15,000bp) and the specific haplogroup of exercise participants. For instance, 3’UTR (untranslated regions) variants that do not directly affect protein function may however affect translation, mRNA shuttling to specific organelles, or epigenetic modification such as microRNA silencing (21). Intronic variants may also lead to splice site changes directly contributing altered protein structure and function (22). As Next Generation Sequencing has become more widely available and affordable, sequencing of the whole mitochondrial genome (16,569 bp) is now feasible to uncover genetic variants associated with physical fitness phenotypes. When used in combination with SNP genotyping arrays, it is possible to examine, not only the 37 mitochondrially-encoded genes, but variants within all nuclear NEMP genes simultaneously.

Therefore, the aim of the present study was to examine the association between genetic variants (i.e. mitochondrial variants and NEMPs), and aerobic fitness measures in the well-characterised Gene SMART cohort. We hypothesise that by utilising whole-mitochondrial sequencing, we will uncover novel genetic variants associated with exercise responses.

## RESULTS

### Exercise responses and Principal Component Analysis (PCA)

Participant characteristics and response to exercise for all phenotypes are detailed in **Table 1**. P-values shown for delta variables are respective of one tail of a paired samples t-test.

**Table 1:**
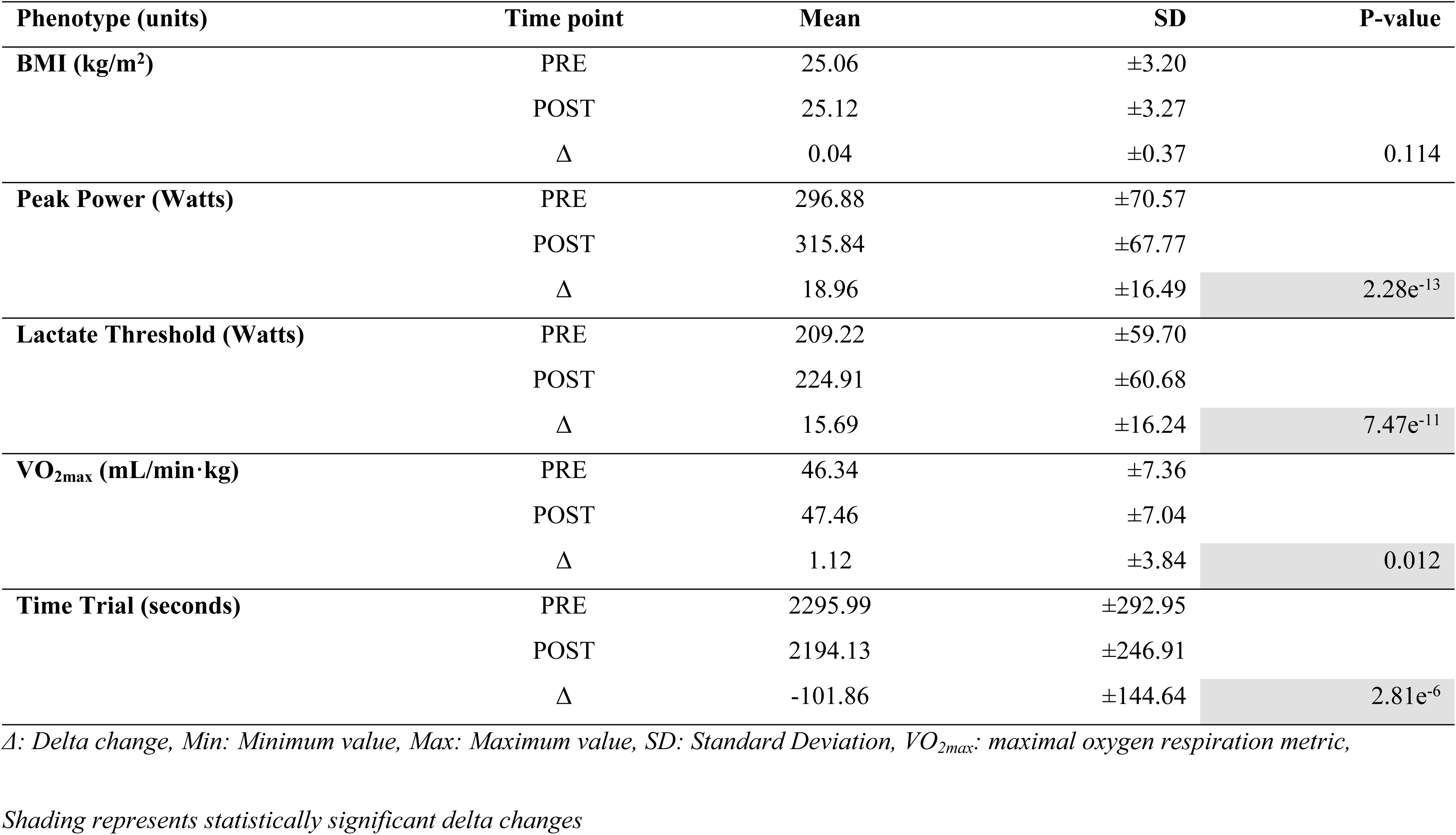
Participant characteristics before and after four weeks of high-intensity interval training in the Gene SMART study.

Four weeks of HIIT elicited small yet significant improvements in Wpeak, LT, VO2max, and TT (PP: 18.96 ± 16.49 Watts, P=2.28e^-13^; LT: 15.69 ± 16.24 Watts, P=7.47e^-11^; VO2max: 1.12 ± 3.84 mL/min**·**kg, P=0.012; TT: −101.86 ± 144.64 seconds, P=2.81e^-6^).

There were 60 distinct haplogroups within the Gene SMART completed cohort of 62 participants. As such, there were no statistically significant associations between the mitochondrial haplogroups with exercise response traits. A summary table of the mitochondrial haplogroups found within the Gene SMART participants is shown in **Table 2**. The confidence scores (0-1) represent the number of mtDNA variants found in each participant that belong to their respective haplogroup.

**Table 2:** Summary of mitochondrial Haplogroups within the Gene SMART study.

| Participant ID | MtDNA Haplogroup | Confidence | Participant ID | MtDNA Haplogroup | Confidence |
| --- | --- | --- | --- | --- | --- |
| SG100 | H1c2a | 0.9505 | SG140 | H1c7 | 0.9581 |
| SG102 | C1b10 | 0.9305 | SG141 | H2a2b3 | 0.9386 |
| SG103 | K1a1b2b | 0.9648 | SG142 | H+152 | 0.8534 |
| SG104 | H6a1b2 | 0.9438 | SG143 | U4a1a | 1 |
| SG105 | H3g | 0.9353 | SG144 | T2b4+152 | 0.9535 |
| SG106 | H94 | 0.8164 | SG145 | H24a | 1 |
| SG107 | K1a1b1a | 0.968 | SG146 | U5b3e | 0.9818 |
| SG108 | J1c3g | 0.9366 | SG147 | U5a1a1 | 1 |
| SG109 | W3a1c | 0.9804 | SG148 | I1a1e | 0.9762 |
| SG110 | H1e1a3 | 0.9486 | SG149 | H6a1a3 | 0.9958 |
| SG111 | H1t | 0.9336 | SG150 | HV | 0.7231 |
| SG112 | K1a4f1 | 0.9641 | SG151 | U5a2b4 | 0.9481 |
| SG113 | T2b+152 | 0.9795 | SG152 | J1c2f | 0.9805 |
| SG114 | U5b1b1+@16192 | 0.9924 | SG153 | K1a4a1 | 0.9783 |
| SG115 | T2b13a | 0.9827 | SG154 | U2e1a1 | 0.94 |
| SG116 | J1c3g | 0.9639 | SG155 | H1a1 | 0.9505 |
| SG117 | H10 | 0.9356 | SG156 | H1a | 0.9898 |
| SG118 | H16b | 1 | SG157 | H3u1 | 0.8918 |
| SG119 | U3a1c1 | 0.9499 | SG158 | H1e1a2 | 0.9243 |
| SG120 | T2b1 | 0.9904 | SG159 | U4b1a2 | 0.9924 |
| SG121 | H15a1a1 | 0.9175 | SG160 | U8a1a | 0.9319 |
| SG122 | K1a | 0.9508 | SG161 | K1a | 0.9204 |
| SG123 | K1a4a1a+195 | 0.9941 | SG162 | U4b1a2 | 0.9924 |
| SG124 | H3 | 0.9852 | SG163 | H4a1a2a | 0.9818 |
| SG125 | L0d2a1a | 0.9839 | SG164 | H2a2b4 | 0.9037 |
| SG126 | H5a1 | 1 | SG165 | T2f1a1 | 0.9306 |
| SG127 | H2b | 0.8848 | SG166 | H1a1 | 1 |
| SG128 | H1 | 0.8676 | SG167 | U5a1b1d+16093 | 0.9791 |
| SG129 | H24a2 | 0.9202 | SG168 | H2a2a1 | 0.5 |
| SG130 | J2a1a1 | 0.9726 | SG169 | T2b | 0.9918 |
| SG131 | U8b1a1 | 0.9258 | SG170 | J1b1a1a | 0.9857 |
| SG132 | V10a | 0.9673 | SG171 | H6a1b3 | 0.985 |
| SG133 | HV1a1a | 0.9296 | SG172 | I2 | 0.9222 |
| SG134 | J1c7a | 0.9841 | SG173 | I2c | 0.9577 |
| SG135 | R1a1a2 | 0.9875 | SG174 | M1a | 0.905 |
| SG136 | HV6a | 0.951 | SG175 | W5 | 0.9513 |
| SG137 | H2a1e1a | 0.9591 | SG176 | T2f1a1 | 0.8887 |
| SG138 | H1b1+16362 | 1 | SG177 | K1a16 | 0.9932 |
| SG139 | J2b1a2a | 0.9655 |  |  |  |

Following PCA on the response traits, we found that the first 4 principal components (PC1: 35.49%, PC2: 28.46%, PC3: 16.51%, PC4: 12.74%) cumulatively explained 92.3% of the total variance between individuals; therefore we included only these first 4 PCs in subsequent analyses.

Following quality control, 170 mitochondrial and 4,124 NEMP genetic variants were included in association testing. A cumulative total of 4,325 NEMP variants and 28 mitochondrial variants passed the nominal threshold of significance (P_unadjusted_ <0.05) for all tests. A solar plot showing the clustering of mitochondrial genomic variants for each trait is shown in **Fig 1** (23)(23)(33). The exonic variants passing the nominal threshold from the mitochondrial association results are summarised in **Table 3**.

**Table 3:** Exonic mitochondrial SNPs associated with phenotypic traits and PCs.

| Trait | CHR | SNP | Allele | Gene | Consequence | Model | MAF | SE (95% CI) | P-value* | FDR | Effect size (beta) |
| --- | --- | --- | --- | --- | --- | --- | --- | --- | --- | --- | --- |
| Δ-LT | MT | 8701 | G | ATP6 | Missense | ADD | 0.032 | 11.24 (-48.2 - -4.2) | 0.023 | 0.39 | -26.19 |
|  | MT | 10873 | C | ND4 | Synonymous | ADD | 0.032 | 11.24 (-48.2 - -4.2) | 0.023 | 0.39 | -26.19 |
|  | MT | 12705 | T | ND5 | Synonymous | ADD | 0.081 | 7.25 (-29 - -0.5) | 0.046 | 0.49 | -14.75 |
|  | MT | 15043 | A | CYB | Synonymous | ADD | 0.113 | 6.08 (-29 - -5.2) | 0.0067 | 0.28 | -17.10 |
| PC3 | MT | 10873 | C | ND4 | Synonymous | ADD | 0.032 | 0.72 (0.17 – 3.0) | 0.032 | 0.54 | 1.57 |
|  | MT | 8701 | G | ATP6 | Missense | ADD | 0.032 | 0.72 (0.17 – 3.0) | 0.032 | 0.54 | 1.57 |
| PC4 | MT | 11467 | G | ND4 | Synonymous | ADD | 0.258 | 0.25 (-1.0 - -0.03) | 0.041 | 0.69 | -0.53 |
|  | MT | 12308 | G | tRNA<sup>Leu</sup> | - | ADD | 0.258 | 0.25 (-1.0 - -0.03) | 0.041 | 0.69 | -0.53 |
|  | MT | 12372 | A | ND5 | Synonymous | ADD | 0.258 | 0.25 (-1.0 - -0.03) | 0.041 | 0.69 | -0.53 |

*CHR: Chromosome #, SNP: Single Nucleotide Polymorphism, MAF: Minor Allele Frequency, SE: Standard Error, CI: Confidence Interval, FDR = False Discovery Rate, $\Delta$ :* *delta change, ADD: Additive model*
*\*P-value adjusted for age*

**Fig 1:**
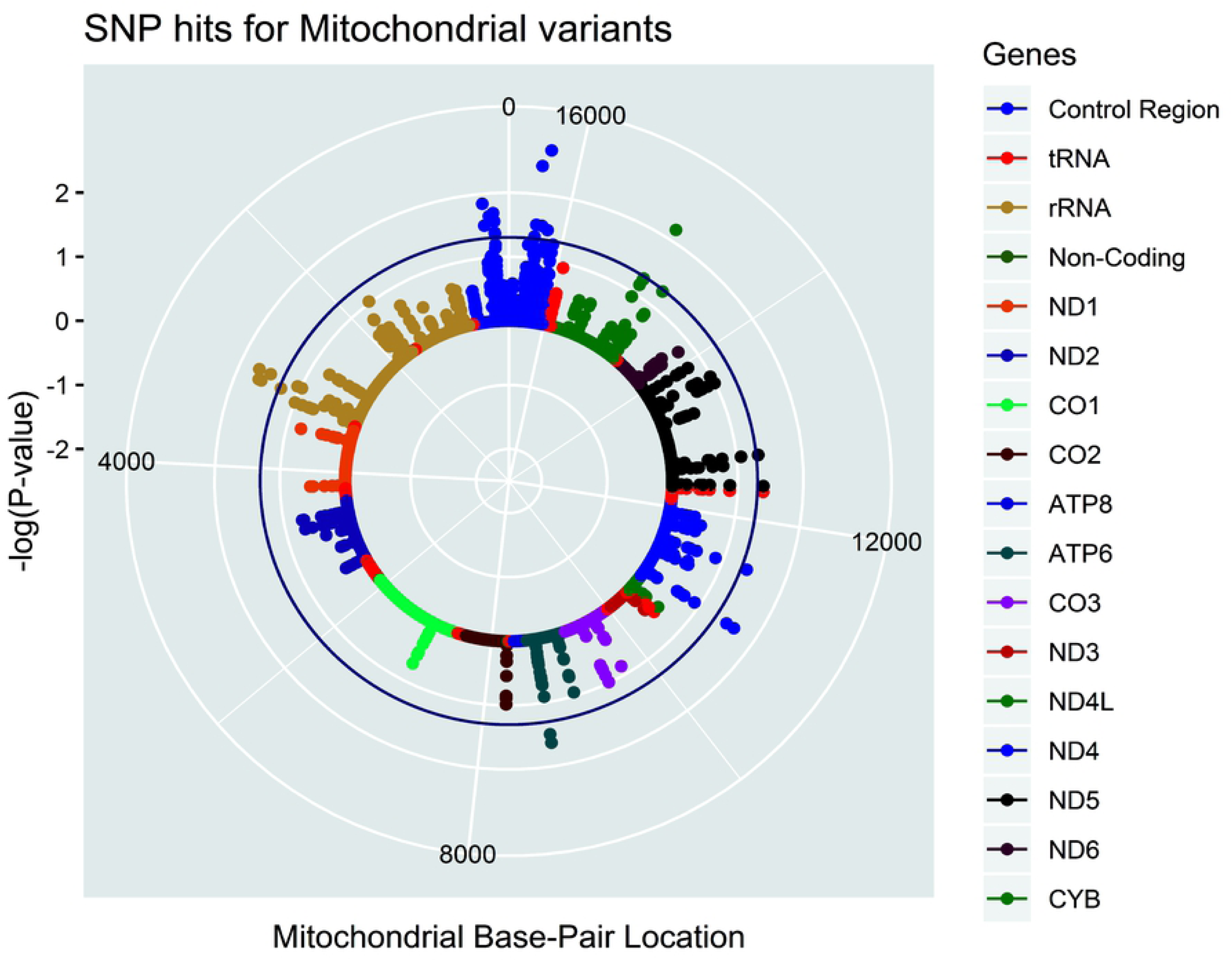
Solar plot showing significant hits from mitochondrial association testing. Each dot represents a detected variant. The inner ring of the plot represents the mitochondrial genome and is coloured based on genomic region as summarised in the plot legend. The X-axis represents the mitochondrial base pair location. The Y-axis represents the significance level [-log10 (P-value)] in the Gene SMART population over multiple traits. The significance threshold was set at P<0.05 and is represented by the circular blue line. The concentric white rings surrounding the genome represent the P-value thresholds −log10 (0.01) and −log10 (0.001) respectively.

28 variants passed the nominal significance threshold of P_unadjusted_ <0.05 in various delta traits and principal components. Of these, 8 were located within the hypervariable control region and therefore discounted from further analyses. A further 2 genetic variants were located within a mitochondrial rRNA gene, 1 within the *tRNA*^*Leu*^ gene, 1 within the *mitochondrially encoded ATP synthase membrane subunit 6* (*ATP6)* gene, 2 within the *mitochondrially encoded NADH: ubiquinone oxidoreductase core subunit 4* (*ND4)*, 2 in *mitochondrially encoded NADH: ubiquinone oxidoreductase core subunit 5* (*ND5)* and 1 in mitochondrially encoded cytochrome B (*CYB)*. None of the mitochondrial genomic variants were associated with composite response traits or individual response traits at FDR < 0.05. A manhattan plot of the NEMP variants is shown in **Figure 2**. A summary of the association statistics for the variants passing a nominal threshold of P_unadjusted_ <1e^-4^ in both the NEMP associations are shown in **Table 4**.

**Table 4:**
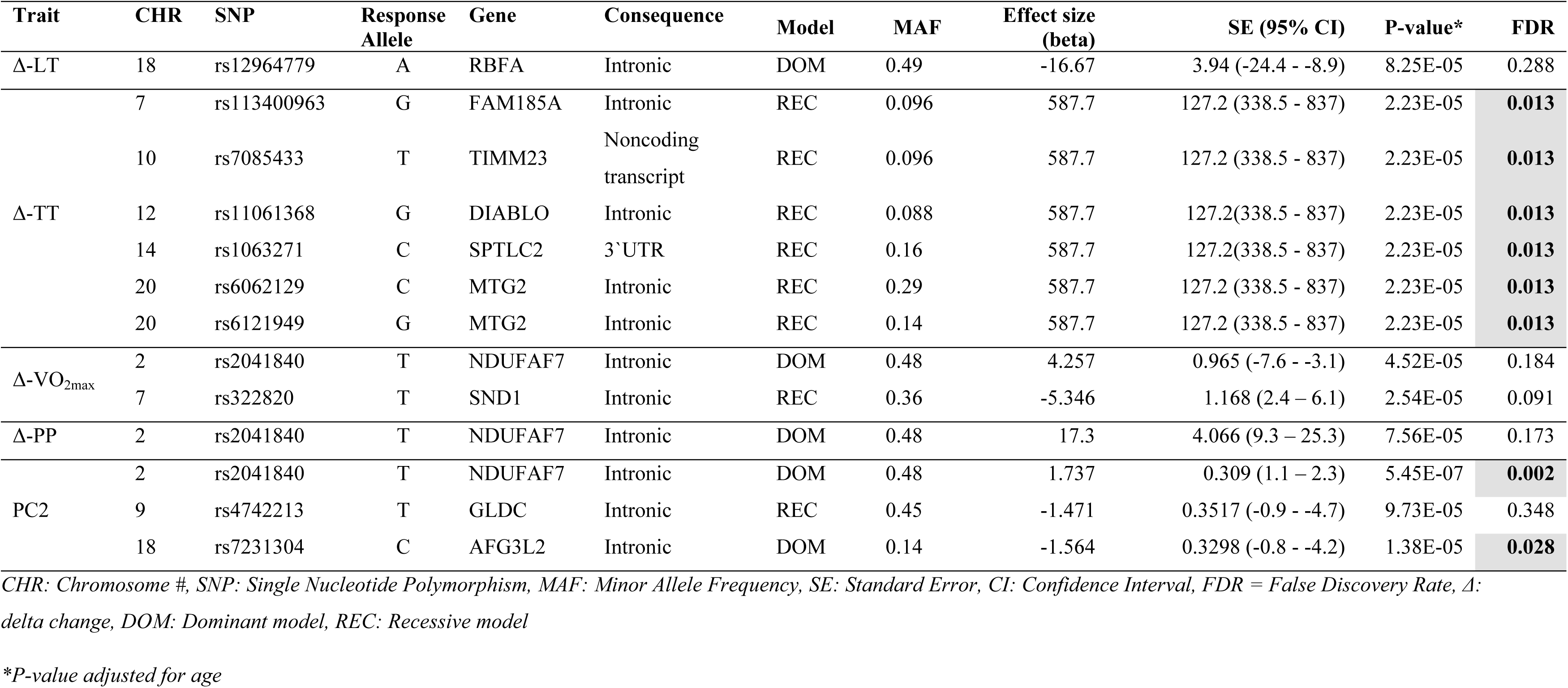
Summary statistics for exonic variants in the nuclear encoded, mitochondria-related genes.

**Fig 2:**
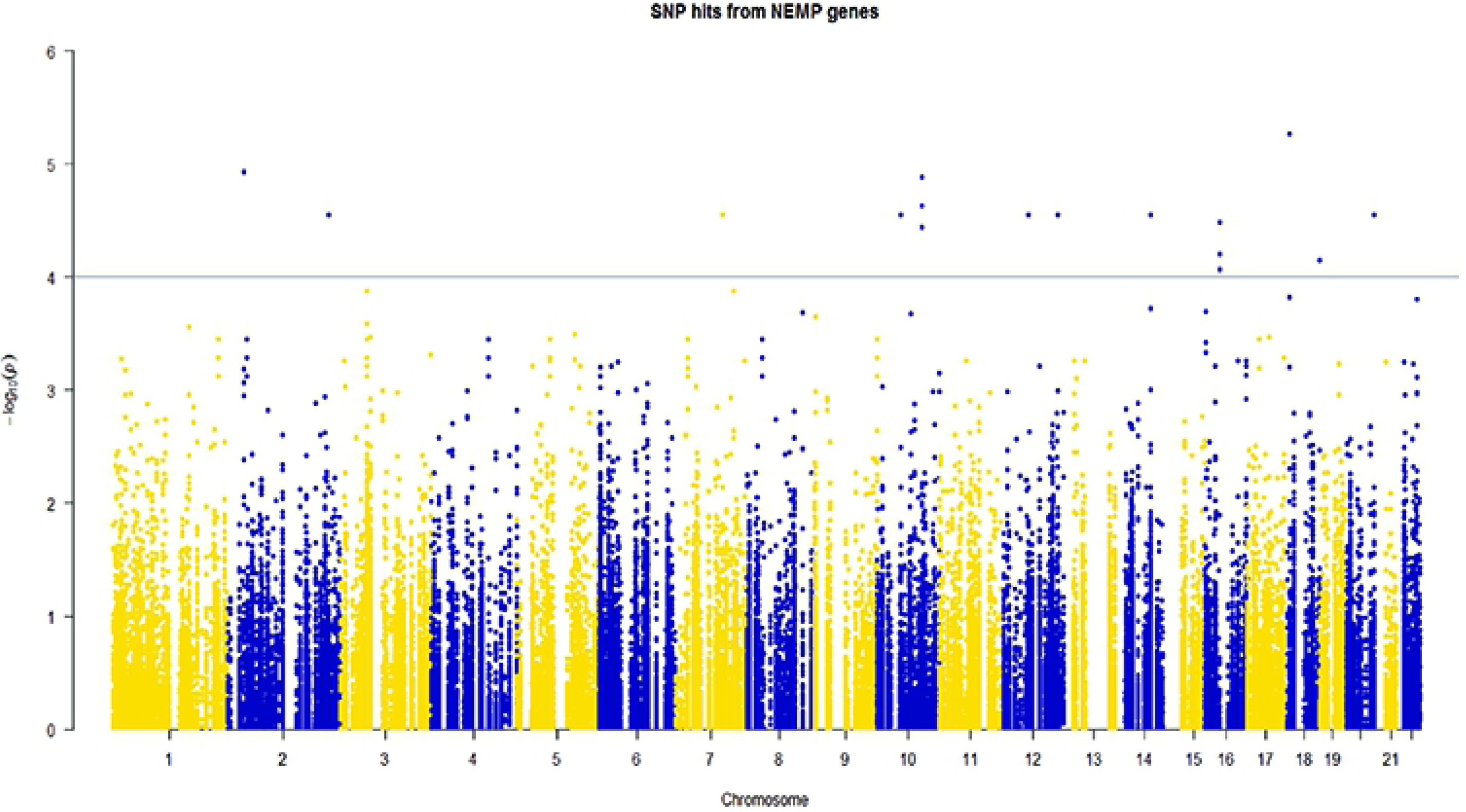
Manhattan plot for all hits from all response phenotypes, biochemical measures, and PCs in the linear dominant and recessive association models. Suggestive significance was set at –log10(P_unadjusted_ = 0.0001, blue line). As all traits were included clusters of variants represent association across multiple traits rather than one significant locus commonly associated with GWAS.

A full list of variants reaching the nominal P value threshold (<0.05) may be found in (**Supplementary Table SI**). 6 SNPs in 5 distinct genes were associated with ΔTT and 2 SNPs in 2 distinct genes were associated with PC2. The most significant variant was rs2041840 associated with PC2 and located within *NDUFAF7;* we found that the rs12712528 variant also within *NDUFAF7* had a moderate correlation with rs2041840 (R^2^=0.5) **Figure 3a**. This variant was also trending towards significance in the Δ-Weight and Δ-VO_2max_ response phenotypes (**Table 3**). The T allele at rs2041840 was associated with a better response to exercise. The Locus Zoom plot (**Fig 3b**) surrounding the *MTG2* gene was also gene-rich with 11 proximal genes. The two associated variants (rs6062129 and rs6121949) were moderately correlated (R^2^=0.5), however there were no SNPs found within the proximal genes. The locus zoom plot for the variants found within the *AFG3L2* gene (**Fig 3c**) was proximal to 6 genes within 200Kb. There was also a proximal SNP within the *SLMO1* gene however this was not in linkage with the variants identified within the *AFG3L2* gene.

**Fig 3.**
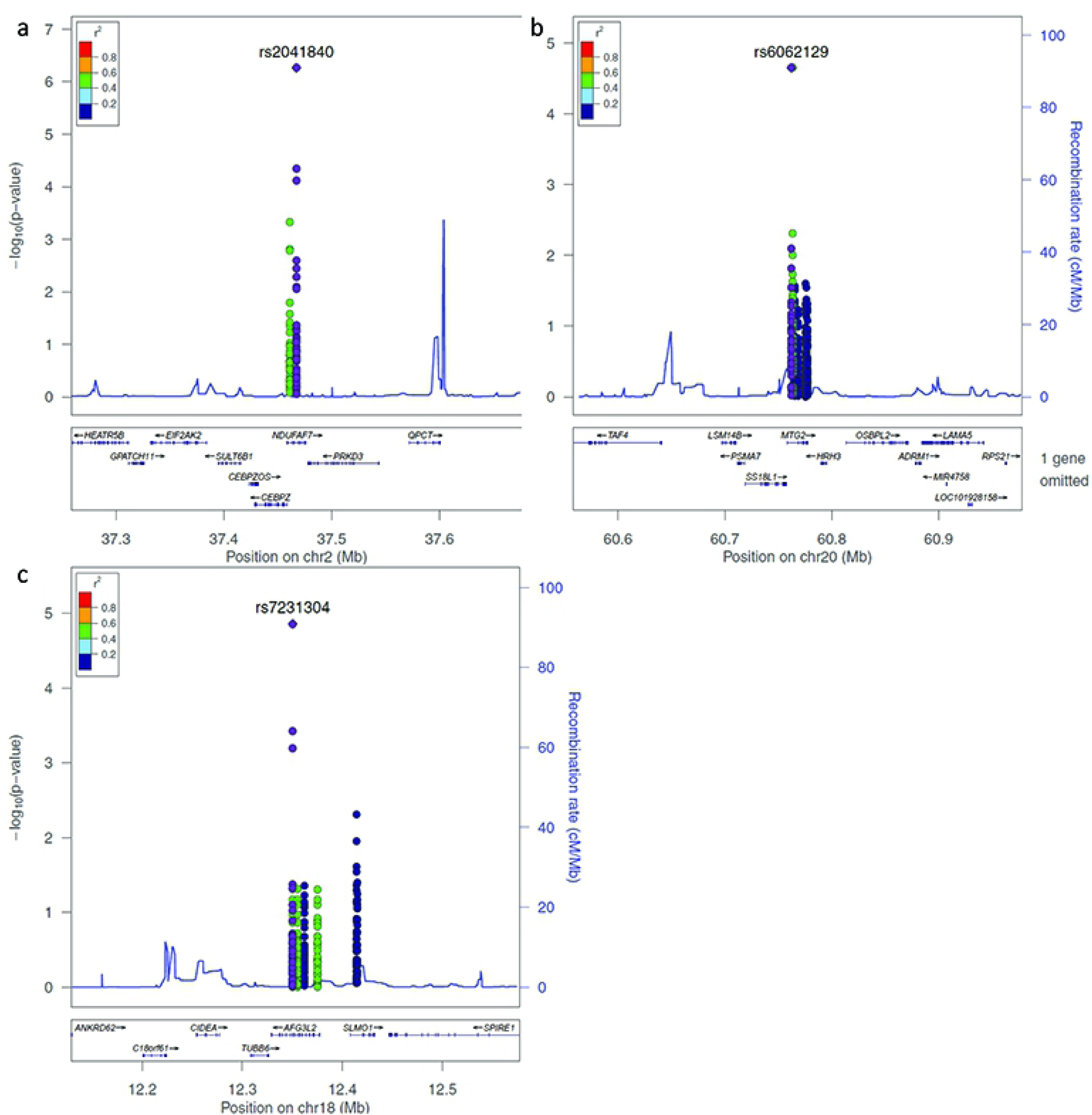
Locus Zoom plots of significant intronic SNPs from the nuclear mitochondrial association testing. Each panel shows the locus surrounding a) rs2041840 within the NDUFAF7 gene, b) rs6062129 variant within the MTG2 gene, and c) rs7231304 variant within the AFG3L2 gene. All panels show the gene of interest ±200Kb. Left y-axis shows –log10(p-value) of association results for all traits and right y-axis shows recombination rate across the locus in relation to the variant of interest. X-axis shows genomic position across the respective chromosomal regions.

## DISCUSSION

In this study, we utilised state-of-the-art mitochondrial sequencing, along with high-throughput targeted genotyping of mitochondrial-related variants encoded by the nucleus (NEMPs) to discover novel genetic variants associated with responses to exercise. A total of 28 mitochondrial and 4,325 nuclear encoded mitochondrial associated variants passed the nominal significance thresholds for the various candidate gene association tests. We did not detect mitochondrial variants associated with exercise response, but we uncovered eight NEMPs in seven distinct genes associated with exercise response.

### Novel exercise loci

The most significant variant was associated with the composite exercise response phenotype and located within an intron of *NDUFAF7* (rs2041840). The T allele was associated with better exercise response as shown by the positive beta values. *NDUFAF7* encodes an arginine methyltransferase that is essential for mitochondrial complex I assembly (24). We have showed that this variant was in a gene rich region with 8 proximal genes (**Fig 3a**), indicating possible effects for this variant in any of the proximal genes or indeed for genes that may be further away from the loci.

The two intronic variants within the *MTG2* gene were found to be associated with the change in time trial measures and appeared to be moderately linked (**Figure 3b**). The *MTG2* gene resides in a gene rich locus with 11 proximal genes. The MTG protein regulates the assembly and function of the mitochondrial ribosome. As such, dysregulation of the gene could result in the downregulation of mitochondrial transcription, and therefore a lower response to exercise training. The variants also showed a 20% recombination rate with the 5’ region of the *TAF4* gene. The TAF4 protein forms part of the transcription factor II D (TFIID) complex and has a central role in mediating promoter responses to transcriptional activators and repressors. Dysregulation of this gene could introduce global translational repression and therefore lack of response to HIIT training. This is supported by the positive effect size for the C and G alleles of the MTG2 variants rs6062129 and rs6121949 respectively (β = 587.7 seconds).

An intronic variant within *AFG3L2* was also shown to be associated with the composite exercise response phenotype (rs7231304), but this gene has not previously been associated with exercise response. However, mutations in *AFG3L2* have been shown to cause spinocerebellar ataxia through the development of mitochondrial proteotoxicity (25, 26). As such, the intronic variation within this gene might inhibit exercise response through dysregulation of mitochondrial structure and function. Further, this variant is in a locus with 6 proximal genes (**Fig 3c**), however no genes within this locus shared a recombination rate above 10%. There were two proximal SNPs with a moderate correlation to the SNP of interest also within the *AFG3L2* genic region.

The T allele at the exonic rs7085433 variant in the *TIMM23* gene was associated with the change in time trial phenotype (Δ-TT) causes a non-coding transcript of the *TIMM23* gene. This gene is one of the targets of transcriptional activators NRF-1 and GA binding protein (GABP/NRF-2) (27), in which we have previously shown genetic variants associated with athletic performance (28, 29). TIMM23 is one of the mitochondrial transmembrane subunits that form the mitochondrial protein import (TIM23) complex. Therefore, this subunit is essential for the transport of peptide containing proteins across the inner mitochondrial membrane. The non-coding transcript resulting from the variant would render the complex non-functional and as such impaired transport of biomolecules across the inner mitochondrial membrane may impair exercise potential. The effect size of this variant was very highly positive (β = 587.7 seconds) and therefore, this non-coding transcript may result in a slower time to complete the time trial.

The rs1063271 variant lies within the 3’ Untranslated Region (UTR) of the *SPTLC2* gene. UTR variants have been shown to influence transcript half-life; through the dysregulated binding of transcript shuttle proteins; or change the binding site of miRNAs resulting in epigenetic silencing of the gene (30). The SPTLC2 protein is involved in the de novo biosynthesis of sphingolipids by forming a complex with its counterpart; SPTLC1 (31). Overexpression of this protein has also been shown to cause elevated sphingolipid formation and therefore mitochondrial autophagy (32). Much like the *TIMM23* rs7085433 variant, the effect size for time to completion in Time Trial (β = 587.7 seconds) indicated that carriers of T allele/genotype have slower TT and therefore poorer response to exercise when compared to carriers of the C allele/genotype. We hypothesise that the C allele for this variant may induce a novel miRNA binding site in the transcript resulting in the silencing of the SPTLC2 gene.

The rs11061368 variant lies within an intronic region of the *DIABLO* gene. The protein encoded by this gene functions to induce apoptotic processes through the activation of caspases in the Cytochrome C/Apaf-1/caspase-9 pathway. We hypothesise that the dysregulation of the *DIABLO* gene could prevent adequate muscle remodelling resulting in the lack of response to training. The variant also lies ∼50Kb away from the *Interleukin 31* (*IL31)* gene, a pro-inflammatory cytokine associated with the activation of Signal Transducer and Activator of Transcription 3 (STAT3) pathways.

### Mitochondrial

None of the mitochondrial genetic variants identified in this study were associated with exercise response in the present study to a threshold of FDR<0.05. Additionally, we lacked enough statistical power to associate mitochondrial haplogroup with exercise responses as the cohort was extremely heterogenous.

The g.A8701G variant within the *ATP6* gene causes a missense change within its respective protein (p.Thr59Ala) and has been well characterised in hypertensive cases (33). This variant was nominally significant in both the Δ-LT phenotype and the PC3 composite trait within the cohort. As the Δ-LT trait was provides a smaller contribution to PC3, the variant was assumed to be partially associated with a mixture of the Δ-TT and Δ-VO_2max_ phenotypes. The effect size of this variant indicated a poor response to exercise training (β = −26.19 LT).

Interestingly, all the variants associated with PC4 were related to the utilisation of the amino acid Leucine. Firstly, the g.A12308G variant within the mitochondrial coding region for the tRNA for Leucine. Whilst the effect of this variant was unclear, it appears to have influenced the composite phenotypes within PC4. Mutations within tRNA genes have previously been associated with reduction in organelle quantity and downregulation of protein synthesis (34). Secondly, both synonymous variants in the ND4 (g.A11467G) and ND5 (g.G12372A) genes result in a codon that is used far less frequently (CUA_[276]_ > CUG_[42]_) in mitochondrial translation processes (35). As the biosynthesis of tRNAs is costly with respect to intracellular energy levels, it is possible that the combination of the dysregulation of the tRNA^leu^ and the codon usage frequency change in two subunits of the mitochondrial membrane respiratory chain NADH dehydrogenase (complex I) may result in premature intracellular energy (ATP) deficiency and contribute to the poor response to exercise training associated with these traits. It should be noted that the stringent thresholds for association in the mitochondrial association tests could also have resulted in false negative results. Additionally, mitochondrial genetic variants rarely influence phenotypic traits in isolation.

We have identified novel nuclear-encoded, mitochondrial-related SNPs and loci associated with adaptations to High Intensity Interval Training. Additionally, we have postulated the mode of action for different molecular mechanisms that may be responsible for the variability in response to exercise intervention. It should be noted that performing mitochondrial sequencing on muscle tissue as opposed to blood may yield more informative results with heteroplasmic associations due to the high concentration of mitochondria. We note that while we have utilised comprehensive sequencing and high throughput arrays in combination with robust exercise phenotypes, the variants associated with responses in this study, need to be replicated in larger cohorts of both the general population and elite athletes. This could be achieved by leveraging on large multi-centre initiatives such as the Athlome consortium (36). Additionally, functional genomic analyses are required to determine the effect of these variants on the molecular pathways commonly involved in exercise response. Such studies could include transcriptomics, epigenetics and functional cell work in a multi-omics approach.

## MATERIALS AND METHODS

### Participants

At the time of analysis, 77 participants had taken part in the study, 62 of whom successfully completed 4 weeks of High-Intensity Interval Training (HIIT) intervention protocol in the Gene SMART (Skeletal Muscle Adaptive Response to Training) study (37) at Victoria University, Australia. Ethical clearance for this study was provided by the Human Research Ethics Committee at Victoria University (Approval Number: HRE13-233), and the clearance was transferred to and also provided by the QUT Human Research Ethics Committee (Approval Number: 1600000342). We analysed the 62 participants who did not drop out of the study and all had healthy BMI and were moderately trained with an age range of (31.33 ± 7.94 years).

The Gene SMART study design has been previously reported (37). Briefly, participants were required to provide medical clearance to satisfy the inclusion criteria. Following familiarisation, baseline exercise performance was determined on a cycle ergometer during a 20 km time trial (TT), and two graded exercise tests (GXTs); these tests were administered a few days apart and no more than two weeks apart to limit temporal variability in performance.

### Molecular Methods

Genomic DNA was extracted for 77 participants regardless of completion status from 2.0mL of whole blood using a QIAmp DNA blood midi kit (QIAGEN, Hilden, Germany). Briefly, the concentration and purity of genomic DNA (gDNA) from all samples was assessed via Nanodrop spectrophotometry and Qubit fluorometry. We used an in-house sequencing method recently developed by our group at the Genomics Research Centre, Queensland University of Technology, Australia to sequence the whole mitochondrial genome of each participant (38). Illumina Infinium Microarray was used on HumanCoreExome-24v1.1 bead chip to genotype all samples for ∼550,000 loci. For all samples, 1µg total gDNA was sent to the Australian Translational Genomics Centre, Queensland University of Technology Australia, for SNP genotyping on the arrays.

### Data Filtering

A bioinformatics pipeline (*SAMtools, BCFtools*) was utilised to generate variant call files (VCF) for all samples as described previously (38). VCF files were then aligned to the *revised Cambridge Reference Sequence* (rCRS) and all sequences were stringently left aligned back to this reference genome to account for the single end (SE) reads generated from Ion Torrent sequence information. FASTA files were generated for all samples and then merged VCF and FASTA files were produced for the entire data set. The merged FASTA files were annexed using MITOMASTER, a mitochondrial sequence database, to call haplogroups and obtain variant annotation information for all samples (39)(25). The merged VCF file was converted to PLINK (v1.90p) format using the function ‘*--make-bed’* for further association analysis.

The *ped* file generated from Illumina GenomeStudio v2.0 software was converted into binary format. We did not impute any genotypes to prevent false positive associations and a larger multiple testing burden. There were 551,839 typed SNPs; subsequent SNP and individual filtering and trimming was based on **1)** SNPs with > 20% missing data (239 removed), **2)** individuals with > 20% missing data (0 removed), **3)** minor allele frequency < 0.01 to remove rare variant associations (260,269 removed), **4)** SNPs out of Hardy Weinberg equilibrium for quantitative traits (58 removed due to P<1e^-6^) (40). All samples passed kinship and heterozygosity thresholds after the filtering outlined above, leaving 62 samples and 291,273 SNPs to analyse. A BED file containing the genomic locations (GRCh37) of all known Nuclear Encoded Mitochondrial Protein genes (NEMPs) was obtained from the Broad Institute’s human MitoCarta2.0 website (41-44). PLINK was used to extract the SNPs within the genomic locations from the Omni Express SNP chip data of the same participants. In total, 4,806 SNPs were within the NEMP genomic regions detailed by the Broad Institute MitoCarta2.0 bed file and considered to be mitochondrially related variants.

### Exercise-response phenotypes

Participant stratification into high and low response groups lead to a loss of statistical power in association testing. As such, and to avoid classifying responders and non-responders via arbitrary thresholds, we chose to keep the phenotypes as continuous variables for association testing (45).

To ascertain variants that were associated with exercise response for key physiological traits, we utilised the delta (Δ) change (Post phenotype – Pre phenotype) quantitative trait data for; peak power output (ΔWpeak in Watts); power at lactate threshold (ΔLT in Watts); peak oxygen uptake (ΔVO_2peak_ in mL/min/kg body weight); and time to completion measurement for a 20 km time trial (ΔTT in seconds). As the quantitative traits were all continuous and to keep maximal statistical power, we did not use arbitrary response thresholds. With multiple, correlated response phenotypes, we conducted a Principal Component Analysis (PCA) of the response phenotypes using the R package *FactoMineR (46).* PCA is a dimensionality reduction method that computes linear combinations of the multiple response phenotypes into principal components (PCs) so that the variance between individuals is maximised. Every individual is then represented by one value for each PC, considered a composite trait of the different response phenotypes. A more detailed description of PCA for composite trait association testing is shown in **Supplementary Fig 1**.

Missing phenotypic values were excluded from the phenotype table prior to PCA to prevent skewing of data and to maintain appropriate PCs. Following the PCA, these variables were set as “missing” for the association analysis. We also tested the individual response phenotypes and compared the significance levels of variants between the composite traits with those within each PC. This resulted in 4 PCs that cumulatively explained > 90% of the variance between participants.

### Statistical analysis

Analysis of the response traits was performed in SPSS using a paired samples t-test. SPSS was also used to test associations between mitochondrial haplogroups and exercise response with a Wald test. Analyses for the mitochondrial SNPs and NEMP SNPs were kept separate for analysis using different association models. We used PLINK V1.90p to perform quantitative linear association tests (95% CI) with both dominant and recessive models. An additive model was also attempted but yielded the same results as our dominant model. We adjusted all association results for age and effect sizes were determined using raw beta regression coefficient values (i.e. genotype X is associated with β [unit specific to trait of interest] changes in the phenotype). Variants that passed a nominal P value threshold of P<0.05 were considered for further analysis whereas variants that passed multiple testing adjustment using the Benjamini-Hochberg False Discovery Rate (FDR<0.05) method were considered significant associations. We performed adjustment for multiple testing for each phenotype separately. All variants from the association tests were plotted in R using the *tidyverse, ggplot2*, and *qqman* packages.

## Acknowledgements

This research was supported by Commonwealth Collaborative Research Network funding to Bond University CRN-AESS. Mr Nicholas Harvey was supported by a PhD stipend also provided by Bond University CRN-AESS. This research was also supported by infrastructure purchased with Australian Government EIF Super Science Funds as part of the Therapeutic Innovation Australia - Queensland Node project (LRG). Nir Eynon is supported by the National Health and Medical Research Council (NHMRC), Australia (NHMRC CDF# APP1140644). Sarah Voisin is also supported by the NHMRC (ECF# APP1157732) and by the Jack Brockhoff foundation.

---

**Supplementary Table S1: NEMP association results for each phenotype and PC in the gene SMART study.** Output file is a compilation of multiple association results and is shown in Plink format. An additional “trait” column has been added to illustrate which association test the results are from.

**Supplementary Fig 1: Summary figure of composite trait building with a PCA method.** The traits shown are individual colours and contribute different amounts to each principal component from the analysis. Then each PC will correspond to each participant that contributes to the original traits. Therefore association of a PC is equal to association of the contribution to each PC from the participants.

